# Protective efficacy of a SARS-CoV-2 DNA Vaccine in wild-type and immunosuppressed Syrian hamsters

**DOI:** 10.1101/2020.11.10.376905

**Authors:** Rebecca L. Brocato, Steven A. Kwilas, Robert K. Kim, Xiankun Zeng, Lucia M. Principe, Jeffrey M. Smith, Jay W. Hooper

**Affiliations:** Virology Division, United States Army Research Institute of Infectious Diseases, Frederick, MD; Pathology Division, United States Army Research Institute of Infectious Diseases, Frederick, MD

**Keywords:** SARS-CoV-2, DNA vaccine, jet injection, immunosuppressed hamster

## Abstract

A worldwide effort to counter the COVID-19 pandemic has resulted in hundreds of candidate vaccines moving through various stages of research and development, including several vaccines in phase 1, 2 and 3 clinical trials. A relatively small number of these vaccines have been evaluated in SARS-CoV-2 disease models, and fewer in a severe disease model. Here, a SARS-CoV-2 DNA targeting the spike protein and delivered by jet injection, nCoV-S(JET), elicited neutralizing antibodies in hamsters and was protective in both wild-type and transiently immunosuppressed hamster models. This study highlights the DNA vaccine, nCoV-S(JET), we developed has a great potential to move to next stage of preclinical studies, and it also demonstrates that the transiently-immunosuppressed Syrian hamsters, which recapitulate severe and prolonged COVID-19 disease, can be used for preclinical evaluation of the protective efficacy of spike-based COVID-19 vaccine.

## Introduction

The COVID-19 pandemic has necessitated the rapid development of candidate vaccines and treatments targeting the SARS-CoV-2. Infection with SARS-CoV-2 results in either asymptomatic infection or disease ranging from mild to severe respiratory symptoms ^1^. Many factors contribute to the spread of this virus, including a large number of asymptomatic cases ^2^ and transmission prior to the onset of symptoms ^3^. An effective vaccine would be an invaluable medical countermeasure to protect individuals, prevent transmission, and contribute to containing and ultimately ending this pandemic.

According to the World Health Organization, as of 30 September 2020, there were 41 SARS-CoV-2 vaccines in clinical trials (Phases I, II and III) and 151 vaccines in preclinical development ^4^. Of these vaccines in preclinical development several have been tested for immunogenicity in mice and nonhuman primates. Few have been tested in disease models such as the Syrian hamster model. The Syrian hamster has become a leading animal model for SARS-CoV-2 medical countermeasure testing because it does not require a modified virus, or animal, and there are several similarities to human COVID-19 disease including rapid breathing, lethargy, ruffled fur and moderate (<10%) weight loss ^5 6^. Histopathology includes areas of lung consolidation, followed by pneumocyte hyperplasia as the virus is cleared. At least three candidate vaccines have been tested for efficacy in the Syrian hamster model ^7-9^.

We have developed a Syrian hamster model of severe COVID-19 disease by using cyclophosphamide (CyP) to transiently immunosuppress the hamsters ^10^. In this model, lymphopenia is induced by CyP treatment starting 3 days before exposure to virus. After a relatively low dose of virus (1,000 PFU), the immunosuppressed hamsters develop a protracted disease with >15% weight loss over several days and other indicators of severe disease including high levels of virus in the lungs. Herein, we describe the testing of a jet-injected SARS-CoV-2 DNA vaccine in both wild-type and transiently-immunosuppressed hamsters. Hantavirus DNA vaccines administered at a dosage of 0.2 mg are highly immunogenic in hamsters when administered using jet injection ^11^. Therefore, as an initial proof-of-concept, we opted to use the 0.2 mg dose.

## Results

### Evaluation of DNA vaccine in wild-type hamster model of COVID-19 disease

A SARS-CoV-2 spike-based DNA vaccine, nCoV-S(JET), was constructed by cloning a human-codon-optimized gene encoding the full-length spike protein into a plasmid vector as described in Methods. The plasmid backbone used for this vaccine, pWRG, has been used for hantavirus DNA vaccines that are currently in phase 1 and 2 clinical trials ^12^. Expression of the spike protein from the nCoV-S(JET) was confirmed to express in cell culture (**Suppl. Fig. S1**). In the first vaccine efficacy experiment, groups of 8 hamsters were vaccinated on week 0 and 3 with either 0.2 mg nCoV-S(JET), or 0.2 mg of a MERS-CoV DNA vaccine, or PBS using jet injection (**Fig. 1A**). Sera were collected after 1 vaccination (Wk 3) or 2 vaccinations (Wk 5) and evaluated in a SARS-CoV-2 plaque reduction neutralization test (PRNT) and pseudovirion neutralization assay (PsVNA). SARS-CoV-2 neutralizing antibodies were detected in all of the animals by both assays after the boost (p=0.0156 (PRNT50), p=0.0078 (PsVNA50), Wilcoxon matched-pairs signed rank test, **Fig. 1B**; PRNT80 and PsVNA80 titers shown in **Suppl. Fig. 2A,B)**. Results from the PRNT and PsVNA were acceptably similar (**Suppl. Fig. 3**). The MERS DNA vaccine did not elicit SARS-CoV-2 cross-neutralizing antibodies as measured by PRNT or PsVNA, but all of animals vaccinate with that vaccine developed MERS virus neutralizing antibodies as measured by PsVNA (**Suppl. Fig. 4**).

**Fig. 1.**
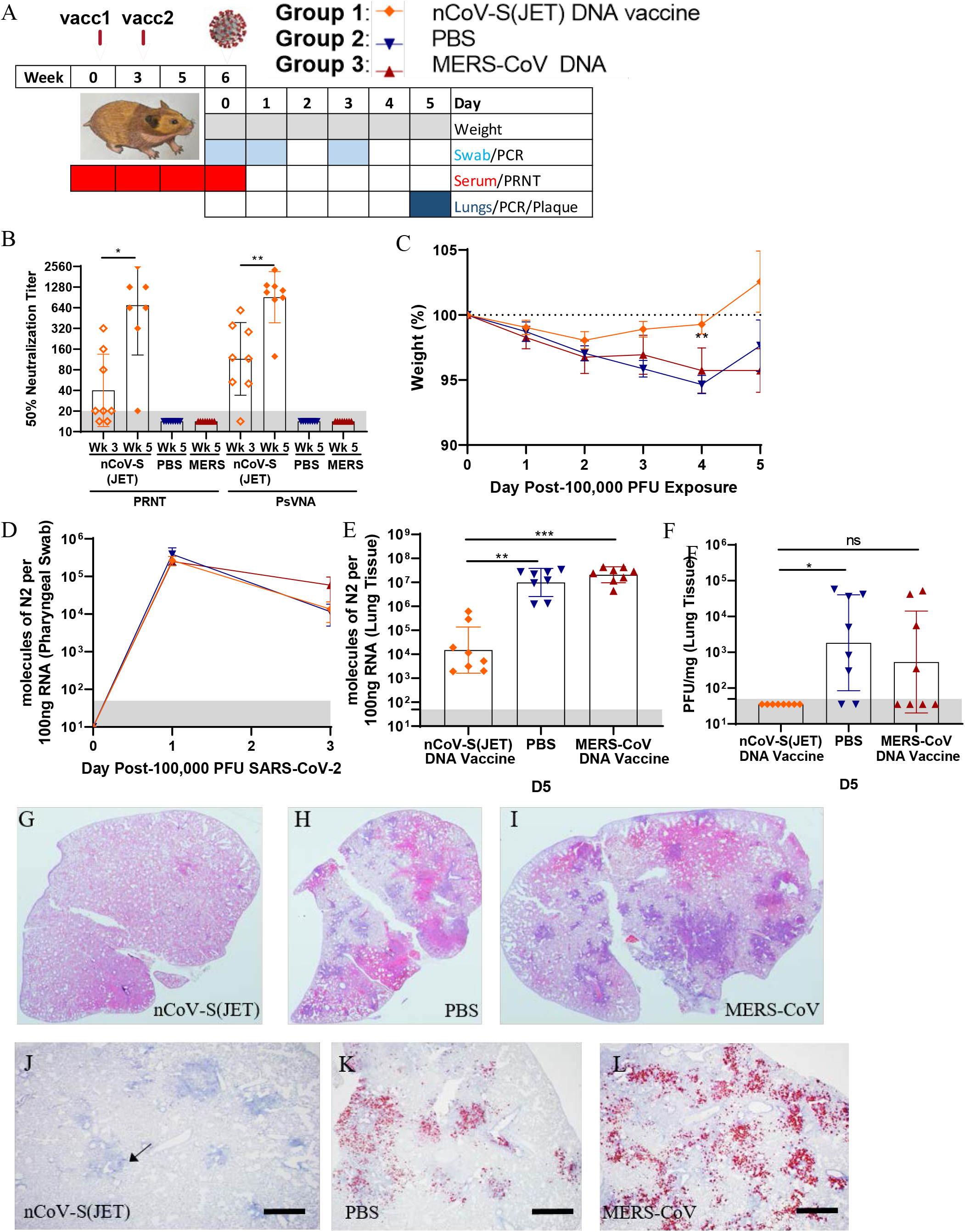
Evaluation of nCoV-S(JET) DNA vaccine in Syrian hamsters. **A)** Experimental design. Groups of 8 hamsters each were vaccinated (vacc) with the nCoV-S(JET) DNA vaccine, PBS, or a MERS-CoV DNA vaccine and then challenged with 100,000 PFU of SARS-CoV-2 virus by the intranasal route. **B)** PRNT50 and PsVNA50 titers from serum collected at indicated timepoints after 1 (open symbols) and 2 (closed symbols) vaccinations (LLOQ = 20, grey shade). **C)** Average animal weights relative to starting weight. Viral RNA in **D)** pharyngeal swabs and **E)** lung homogenates (LLOQ = 50 copies, grey shade). **F)** Infectious virus as measured by plaque assay (LLOD = 50 PFU, grey shade). Bright field imagery of H&E staining of lung sections from **G)** nCoV-S(JET) DNA, **H)** PBS, or **I)** MERS-CoV vaccinated hamsters where purple indicates areas of consolidation. ISH to detect SARS-CoV-2 genomic RNA in lung sections of **J)** nCoV-S(JET) DNA, **K)** PBS, and **L)** MERS-CoV vaccinated hamsters. Rare, positive labeling in nCoV-S(JET) DNA vaccinated hamster lung sections were detected (arrows). Asterisks indicate that results were statistically significant, as follows: *, *P*<0.05; **, *P*<0.01; ***, *P*<0.001; ns, not significant. Scale bars = 400 microns.

**Fig. 2.**
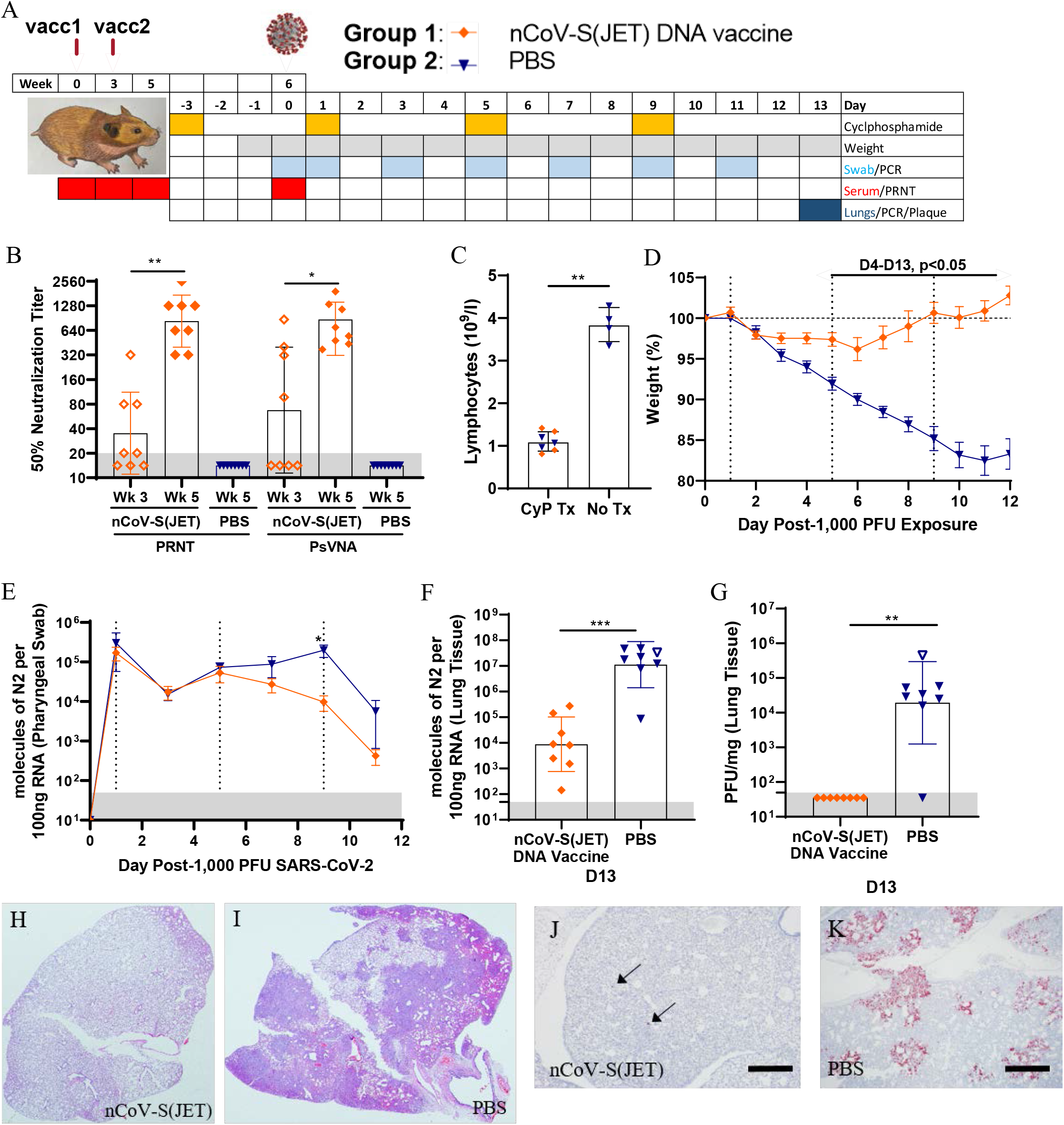
Evaluation of nCoV-S(JET) DNA vaccine in immunosuppressed Syrian hamsters. **A)** Experimental design. Groups of 8 hamsters each were vaccinated (vacc) with the nCoV-S(JET) DNA vaccine or PBS, immunosuppressed with cyclophosphamide (dashed vertical lines in **D)** and **E)**), and then challenged with 1,000 PFU of SARS-CoV-2 virus by the intranasal route. **B)** PRNT50 and PsVNA50 titers from serum collected at indicated timepoints after 1 (open symbols) and 2 (closed symbols) vaccinations (LLOQ = 20, grey shade). **C)** Lymphopenia was confirmed by hematology. **D)** Average animal weights relative to starting weight. Viral RNA in **E)** pharyngeal swabs and **F)** lung homogenates (LLOQ = 50 copies, grey shade). **G)** Infectious virus as measured by plaque assay (LLOD = 50 PFU, grey shade). A single animal from the PBS group succumbed on Day 9 post-exposure (open symbol in **F)** and **G)**). Bright field imagery of H&E staining of lung sections from **H)** nCoV-S(JET) DNA or **I)** PBS vaccinated hamsters where purple indicates areas of consolidation. ISH to detect SARS-CoV-2 genomic RNA in lung sections of **J)** nCoV-S(JET) DNA and **K)** PBS vaccinated hamsters. Rare, positive labeling in nCoV-S(JET) DNA vaccinated hamster lung sections were detected (arrows). Asterisks indicate that results were statistically significant, as follows: *, *P*<0.05; **, *P*<0.01; ***, *P*<0.001; ns, not significant. Scale bars = 400 microns.

Three weeks after the boost all of the hamsters were exposed to 100,000 PFU SARS-CoV-2 by the intranasal route (Day 0). Daily weight change data demonstrated that animals vaccinated with nCoV-S(JET) lost significantly less weight than the animals vaccinated with PBS on Day 4 (p=0.0044, Kruskal-Wallis test, **Fig. 1C**). In contrast, animals vaccinated with the MERS-CoV DNA vaccine were not protected from weight loss. No significant changes in viral RNA load from pharyngeal swabs between nCoV-S(JET)-vaccinated and PBS animals were observed at any timepoint (**Fig. 1D**). Animals were euthanized 5 days after virus exposure and lung homogenates were assayed for viral RNA and infectious virus. There were significant reductions in viral RNA (p=0.0003 vs. MERS-CoV, p=0.0060 vs. PBS, Kruskal-Wallis test with multiple comparisons, **Fig. 1E**) and infectious virus (p=0.0968 vs. MERS-CoV, p=0.0114 vs. PBS, Kruskal-Wallis test with multiple comparisons, **Fig. 1F**) in the hamsters vaccinated with nCoV-S(JET) DNA when compared to the animals vaccinated with MERS-CoV DNA or PBS. Bright field imagery of lung H&E sections show extensive areas of consolidation in PBS- and MERS-CoV-vaccinated hamsters that are not present in nCoV-S(JET)-vaccinated hamsters (**Fig. 1G,H,I**). Multifocal and scattered positive SARS-CoV-2 RNA labeling in areas of inflammation and respiratory epithelial cells is detected by *in situ* hybridization (ISH) in PBS and MERS-CoV vaccinated hamsters that are not present in nCoV-S(JET) vaccinated hamsters (**Fig. 1J,K,L**). Together, these data indicate the nCoV-S(JET) vaccine had a protective effect in the Syrian hamster model.

### Evaluation of DNA vaccine in an immunosuppressed hamster model of severe COVID-19 disease

In a second vaccine efficacy experiment, groups of 8 hamsters were vaccinated on week 0 and 3 with 0.2 mg of nCoV-S(JET) or PBS in a 0.1 mL volume using jet injection (**Fig. 2A**). Sera were collected after 1 vaccination (Wk 3) or 2 vaccinations (Wk 5) and evaluated in a SARS-CoV-2 PRNT and PsVNA (p=0.0078 (PRNT50), p=0.0234 (PsVNA50), Wilcoxon matched pairs signed rank test, **Fig. 2B)**. Correlation of neutralization assays is shown in **Suppl. Fig. 3**. Neutralizing antibodies were detected in all of the animals by both assays after the boost (**Fig. 2B**). In contrast, hamsters vaccinated with PBS had undetectable neutralizing antibodies in both assays. Previously we demonstrated that transient immunosuppression using CyP results in a severe disease model in Syrian hamsters ^10^. Here, hamsters were treated with CyP on Day −3, 1, 5, and 9 relative to challenge. On Day 0 prior to challenge, hamsters were bled for hematology to confirm lymphopenia (**Fig. 2C**). Hamsters were then exposed to 1,000 PFU SARS-CoV-2 by the intranasal route (Day 0). Starting on Day 4 post-infection and continuing through the rest of the experiment, significant differences in weight were observed between hamsters vaccinated with nCoV-S(JET) DNA versus PBS (Day 4, p=0.0059, Days 5-13, p<0.001, t-test, **Fig. 2D**). nCoV-S(JET) DNA-vaccinated hamsters had significantly less viral RNA load in pharyngeal swabs on Day 9 (p=0.0286, t-test, **Fig. 2E**) and trending lower on Days 7 and 11. On Day 13 post-infection (termination of experiment), significant reductions in viral RNA (p=0.0006, t-test, **Fig. 2F**) and infectious virus (p=0.0014, t-test, **Fig. 2G**) in the lungs were measured between the nCoV-S(JET) DNA vaccine and PBS groups. Similar pathology was noted in the transiently immunosuppressed hamsters compared to wild-type animals shown in Figure 1, with extensive areas of consolidation observed by H&E and multifocal and scattered positive SARS-CoV-2 RNA labeling in areas of inflammation and respiratory epithelial cells by ISH (**Fig. 2H,I**). Noteworthy, these lungs were collected on Day 13 whereas those collected in the experiment with wild-type hamsters were collected on Day 5. Together, these data indicate the nCoV-S(JET) vaccine had a protective effect in a SARS-CoV-2 infection model with severe and prolonged disease, in which animals that were transiently immunosuppressed before exposure to virus. Unprotected animals lost >15% of their weight and still harbored infectious virus in their lungs almost two weeks after exposure.

## Discussion

The COVID-19 pandemic has spurred an unprecedented global effort to develop a vaccine to prevent this disease. Nearly every conceivable vaccine platform has been brought to bear on the problem including both RNA- and DNA-based vaccines. Nucleic acid vaccines can be produced rapidly once a target immunogen sequence is known and can be modified rapidly if changes in the sequence become necessary; however, delivering the nucleic acid to cells for immunogen expression remains a technical challenge. For RNA vaccines, efficient vaccine delivery requires formulation with lipid nanoparticles (LNPs) or other modalities to protect the RNA and get it across cell membranes. The safety and efficacy of LNP-formulated RNA is currently being assesses in multiple COVID vaccine trials (Clinicaltrials.gov). DNA delivered by needle and syringe can be immunogenic without LNP formulation, even in nonhuman primates ^13^; however, the use of other techniques such as electroporation or jet injection can increase immunogenicity while reducing dosing requirements. At least one COVID-19 DNA vaccine delivered by electroporation (Inovio) has advanced into the clinic (Clinicaltrials.gov).

To our knowledge, there are no reports of a COVID-19 DNA delivered by jet injection advancing into the clinical- or even progressed to animal efficacy testing. This is surprising because of the logistical and regulatory advantages of disposable syringe jet injection over electroporation. There are several contract manufacturing organizations around the world capable of rapidly producing GMP plasmid for use in humans. Thus, the drug substance could be produced rapidly and the safety profile for DNA vaccines has been established over decades. The drug product delivery system, disposable syringe jet injection, such as PharmaJet’s Stratis, is U.S. FDA 510(k)-cleared and has CE Mark and WHO PQA certification. Disadvantages of the DNA vaccine is that at least one booster vaccination, and possibly two in humans, would likely be needed and the dosage would be milligrams rather than micrograms, as is the case for LNP-formulated mRNA vaccines.

There are a limited number of published reports of COVID-19 vaccine efficacy testing in animal models of COVID-19 disease. These include the testing of self-amplify mRNA in the K18-hACE2 mouse model ^14^, a VSV-vectored vaccine in the hACE2 transduced mouse model ^15^, and at least four virus-vectored (yellow fever, adenovirus, VSV, and inactivated Newcastle disease virus) vaccines in SARS-CoV-2 adapted mouse and/or the Syrian hamster model ^7,9,16,17^. In all of the aforementioned efficacy experiments, the vaccines were based on the full-length spike protein and neutralizing antibodies were predictive of protection. Here we used a used a jet injection technique to deliver a SARS2 spike-based DNA vaccine to Syrian hamsters. Jet injection technology is not widely available for small animal use. We used a human intradermal jet injection technology to deliver vaccines intramuscularly to the hamsters. We had previously demonstrated approximately 300-fold increases in neutralizing antibodies when this jet injection technique was used relative to a needle and syringe in hamsters vaccinated with hantavirus DNA vaccines ^11^.

The immunogenicity parameter we focused on was neutralizing antibody. We measured neutralizing antibodies against live virus by PRNT and a non-replicating VSV-based PsVNA. These assays showed significant correlation (p< 0.0001) (**Suppl. Fig 3A,B**). The neutralizing antibody levels rose significantly after the booster vaccination reaching a PRNT80 geometric mean titer (GMT) of 207 and PRNT50 GMT of 761 that are comparable or exceeding titers of other DNA vaccines evaluated in nonhuman primates ^13^ and mice ^18^. The PRNT50=761 is similar to the 50% titers elicited in hamsters vaccinated with single-dose, live-virus vectored vaccines: Ad26-vectored vaccine PsVNA50 <1000; VSV-vectored vaccine PRNT50 <1000, and Yellow Fever-vectored vaccine PRNT50 <1000 ^7-9^. Neutralization titer was plotted against viral RNA detected in lung tissue collected at the time of euthanasia. Negative correlation was observed (**Suppl. Fig. 5**); however, this did not reach statistical significance. There was no cross-neutralizing antibodies against MERS pseudovirions, and those animals were not protected from disease in the hamster model.

Our results in the transiently-immunosuppressed hamster model add extra credence to the idea that antibodies are playing an important role in the protection observed. The transient-immunosuppressed hamster model is a low-dose (1,000 PFU), severe disease model. CyP treatment, renders B and T cells non-functional, essentially replicating an antibody passive transfer experiment where, rather than the passive transfer of exogenous antibody, the vaccine-generated antibody circulating in the animal prior to CyP treatment must be sufficient to protect against disease. If a candidate vaccine where to protect in the wild-type hamster model but not in the transiently-immunosuppressed hamster model, then that would indicate that T and/or B cell proliferation is required for protection afforded by that vaccine. In the case of the nCoV-S(JET) DNA vaccine, normal T and/or B cell proliferation at the time of exposure was not necessary for the protective effect.

This study shows that a relatively simple unmodified full-length S DNA vaccine administered by a relatively simple jet injection technique can elicit neutralizing antibodies after a single vaccination, PRNT50 titers > 700 after a booster, and protect in two hamster models of disease caused by SARS-CoV-2. The vaccine-mediated protection in the transiently-immunosuppressed hamster model provides additional insights into the mechanism of vaccine-mediated protection against SARS-CoV-2.

## Materials and Methods

### Ethics

Animal research was conducted under an IACUC approved protocol at USAMRIID (USDA Registration Number 51-F-00211728 & OLAW Assurance Number A3473-01) in compliance with the Animal Welfare Act and other federal statutes and regulations relating to animals and experiments involving animals. The facility where this research was conducted is fully accredited by the Association for Assessment and Accreditation of Laboratory Animal Care, International and adheres to principles stated in the Guide for the Care and Use of Laboratory Animals, National Research Council, 2011.

### Plasmid construction

For both pWRG/nCoV-S(opt) and pWRG/MERS-S(opt), the full-length S gene open reading frame, preceded at the N-terminus by Kozak sequence (ggcacc), was human codon usage-optimized and synthesized by Genewiz (South Plainfield, NJ) and cloned into the BglII-NotI site of DNA vaccine vector pWRG. The SARS-nCoV-2 S sequence used was the Wuhan coronavirus 2019 nCoV S gene open reading frame (Genebank accession QHD43416). The MERS sequence used was nc Jordan-N3/2012 S gene open reading frame (Genebank accession AGH58717.1). The plasmids for use in vaccinations were produced commercially and diluted in PBS to 2 mg/mL (Aldevron, Fargo ND). Expression of the spike protein from pWRG/nCoV-S(opt) was confirmed by transfection of 293T cells followed by immunofluoresence antibody test (IFAT) using heat inactivated (56°C 30 min) human convalescent plasma NRS-53265 (ATCC, Manassa, VA) and compared to empty vector (**Suppl. Fig. 1**). A second plasmid for the PsVNA was constructed by deletion of 21 amino acids from the COOH terminus of the full length plasmid, pWRG/CoV-S(opt)Δ21 for better incorporation in to pseudovirions ^19^. The pWRG/nCoV-S(opt) plasmid is also called nCoV-S(JET) when combined with jet injection.

### Animal Vaccinations

Wild type (females only, aged 6-8 weeks) hamsters (*Mesocricetus auratus*) were anesthetized by inhalation of vaporized isoflurane using an IMPAC6 veterinary anesthesia machine. Fur over the semitendinosus and biceps femoris muscles (right leg) were removed using electric clippers. The PharmaJet® Tropis device was used to deliver 0.2 mg of DNA in a 0.1 mL volume intramuscularly ^11^. Specifically, the disposable syringe of the device was pressed against the skin, and the device was activated resulting in the delivery of a liquid jet into the muscle and overlying tissues.

### Other Animal Procedures

In addition to vaccination, the following procedures were conducted after anesthetizing the hamsters as described above: intranasal challenge of virus, cyclophosphamide (CyP) intraperitoneal injections, pharyngeal swabs, and non-terminal blood collection. Intranasal instillation of SARS-CoV-2 was administered in a volume of 50µl for the challenge doses of 1,000 PFU, and 100µl for the challenge dose of 100,000 PFU. CyP treatment (Baxter, pharmaceutical grade) consisted of an initial loading dose of 140mg/kg on Day −3, followed by maintenance doses of 100mg/kg on Days 1, 5, and 9 post-exposure. Pharyngeal swabs in 0.5ml of complete media were used for virus detection to monitor infection and disease course in hamsters. Vena cava blood collection was limited to 7% of total blood volume per week. Terminal blood collection was performed by cardiac injection at the time of euthanasia. All work involving infected animals was performed in an animal biosafety level 3 (ABSL-3) laboratory.

### SARS-CoV-2 stock

An aliquot of the third passage of SARS-CoV-2 USA-WA-1/2020 was received from the CDC and propagated in ATCC Vero 76 cells (99% confluent) in EMEM containing 1% GlutaMAX, 1% NEAA, and 10% heat-inactivated fetal bovine serum at an MOI of 0.01. Supernatant was collected from cultures exhibiting characteristic CPE and clarified by centrifugation (10,000 g x 10 minutes). Clarified virus was subjected to the following specifications: Identification by SARS-CoV-2 RT-PCR assay, Quantification by agarose-based plaque assay, free from contaminants by growth of chocolate agar plates, endotoxin testing using Endosafe® nexgen-PTS, and mycoplasma using MycoAlert test kit, and genomic sequencing. For experiments with a challenge dose of ≤10,000 PFU, virus passage 5 was used; for experiments with a challenge dose of 100,000 PFU, passage 6 was used. Genomic analysis indicates no changes between passage 3, 5, and 6 lots.

### Viral RNA assay

Following 3 freeze/thaws of frozen swabs in media, 250µl of media was removed and added to 750µl of Trizol LS. Approximately 200mg of organ tissue was homogenized in 1.0ml of Trizol using M tubes on the gentleMACS dissociator system on the RNA setting. RNA was extracted from Trizol LS or Trizol per manufacturer’s protocol. A Nanodrop 8000 was used to determine RNA concentration, which was then raised to 100ng/µl in UltraPure distilled water. Samples were run in duplicate on a BioRad CFX thermal cycler using TaqPath 1-step RT-qPCR master mix according to the CDC’s recommended protocol of 25°C for 2 minutes, 50°C for 15 minutes, 95°C for 2 minutes, followed by 45 cycles of, 95°C for 3 seconds and 55°C for 30 seconds. The forward and reverse primer and probe sequences are: 2019-nCoV_N2-F, 5’-TTA CAA ACA TTG GCC GCA AA-3’, 2019-nCoV_N2-R, 5’-GCG CGA CAT TCC GAA GAA-3’, and 2019-nCoV_N2-P, 5’-ACA ATT TCC CCC AGC GCT TCA G-3’. The limit of detection for this assay is 50 copies.

### PRNT

An equal volume of complete media (EMEM containing 10% heat-inactivated FBS, 1% Pen/Strep, 0.1% Gentamycin, 0.2% Fungizone, cEMEM) containing SARS-CoV-2 was combined with 2-fold serial dilutions of cEMEM containing antibody and incubated at 37°C in a 5% CO2 incubator for 1 hour (total volume 222µl). 180 µl per well of the combined virus/antibody mixture was then added to 6-well plates containing 3-day old, ATCC Vero 76 monolayers and allowed to adsorb for 1 hour in a 37°C, 5% CO2 incubator. 3mL per well of agarose overlay (0.6% SeaKem ME agarose, EBME with HEPES, 10% heat-inactivated FBS, 100X NEAA, 1% Pen/Strep, 0.1% Gentamycin and 0.2% Fungizone) was then added and allowed to solidify at room temperature. The plates were placed in a 37°C, 5% CO2 incubator for 2 days and then 2mL per well of agarose overlay containing 5% neutral red and 5% heat-inactivated FBS is added. After 1 additional day in a 37°C, 5% CO2 incubator, plaques were visualized and counted on a light box. PRNT50 and PRNT80 titers are the reciprocal of the highest dilution that results in an 50% and 80% reduction in the number of plaques relative to the number of plaques visualized in the cEMEM alone (no antibody) wells.

### Pseudovirion Neutralization Assay (PsVNA)

The PsVNA used to detect neutralizing antibodies in sera utilized a non-replicating vesicular stomatitis (VSV)-based luciferase expressing system described previously ^20^. For the MERS PsVNA there were no modifications, for SARS-CoV-2 assays there were two modifications: 1) no complement was used to parallel the SARS-CoV-2 PRNT assay, 2) a monoclonal anti-VSV-G (IE9F9) was added at 100ng/ml to eliminate any residual VSV activity in the pseudotype preparation. PsVNA50 and PsVNA80 titers were interpolated from 4-parameter curves, and GMTs were calculated.

### Pseudovirion Production

Pseudovirions were produced using the pWRG/CoV-S(opt)Δ21 or MERS-CoV plasmid described above. HEK293T cells were seeded in T75 tissue culture flasks to be ∼80% confluent the following day and were transfected with the plasmid of interest using Fugene 6 (Promega). After ∼18 h the transfection media was removed and the cells were infected with VSVΔG∗rLuc at a multiplicity of infection of ∼0.07 for 1 h at 37°C. The media was removed and fresh media was added, the flasks were then incubated at 32°C for 72 h. The supernatant from infected cells was collected and clarified by high speed centrifugation, followed by a PEG 8,000 precipitation with 3.2% salt. The PEG mixture is spun at10K xG for 45 min. The pellet was resuspended overnight in 1 mL TNE buffer, then filtered using a 0.45 μm filter, aliquoted and stored at −70°C.

### Plaque Assay

Approximately 200mg of lung tissue was homogenized in 1.0mL of cEMEM using a gentleMACS M tubes and a gentleMACS dissociator on the RNA setting. Tubes were centrifuged to pellet debris and supernatants collected. Ten-fold dilutions of the samples were adsorbed to Vero 76 monolayers (200µl of each dilution per well). Following a 1 hour adsorption in a 37°C, 5% CO2 incubator, cells were overlaid and stained identically as described for PRNT. The limit of detection for this assay is 50 plaque forming units (PFU).

### Hematology

Whole blood collected in EDTA tubes was analyzed on an HM5 hematology analyzer on the DOG2 setting.

### Preparation of tissues for histology

Tissues were fixed in 10% neutral buffered formalin, trimmed, processed, embedded in paraffin, cut at 5 to 6µm, and stained with hematoxylin and eosin (H&E).

### Bright Field Imagery

Photographs of the H&E stained slides were taken with a Canon EOS 7D Mark II (mfr#9128B002AA) and Canon EF 100mm f/2.8L Macro (mfr#3554B002) lens. Slides were placed on a lightbox and photographed at 1:1 magnification with a shutter speed of 1/100sec, aperture of f8.0, ISO 400 and saved as Canon RAW files. Contrast was adjusted equally for all images with Photoshop Lightroom and then exported as PNG files.

### *In situ* hybridization

To detect SARS-CoV-2 genomic RNA in FFPE tissues, *in situ* hybridization (ISH) was performed using the RNAscope 2.5 HD RED kit (Advanced Cell Diagnostics, Newark, CA, USA) as described previously ^21^. Briefly, forty ZZ ISH probes targeting SARS-CoV-2 genomic RNA fragment 21571-25392 (GenBank #LC528233.1) were designed and synthesized by Advanced Cell Diagnostics (#854841). Tissue sections were deparaffinized with xylene, underwent a series of ethanol washes and peroxidase blocking, and were then heated in kit-provided antigen retrieval buffer and digested by kit-provided proteinase. Sections were exposed to ISH target probe pairs and incubated at 40°C in a hybridization oven for 2 h. After rinsing, ISH signal was amplified using kit-provided Pre-amplifier and Amplifier conjugated to alkaline phosphatase and incubated with a Fast Red substrate solution for 10 min at room temperature. Sections were then stained with hematoxylin, air-dried, and cover slipped.

### Statistical analyses

Statistical analyses were completed using GraphPad Prism 8. Weight data was analyzed using a one-way ANOVA with multiple comparisons for experiments with ≥2 groups; unpaired t-tests were used to analyze weight data for experiments with 2 groups. Comparisons of lymphocyte levels and lung viral load was assessed using a one-way ANOVA with multiple comparisons for experiments with ≥2 groups; unpaired t-tests were used to analyze weight data for experiments with 2 groups. Significance of survival data was assessed using log-rank tests. In all analyses, *P*<0.05 is considered statistically significant.

## Data and materials availability

All data is available in the main text.

## Acknowledgments

We would like to acknowledge the Comparative Medicine Division, Histology Lab and Aerobiological Sciences technicians at USAMRIID. We thank Joshua Moore, Jimmy Fiallos, Steven Stephens, Leslie Klosterman, Lynda Miller, Jua Liu, April Babka, Neil Davis and Dave Dyer for assistance with veterinary care, histology and molecular assays, and hematology. Additionally, we thank Brian Kearney, Kathleen Gibson and the Unified Culture Collection for providing the virus.

## Author Contributions

R.L.B. and J.W.H. designed the study; R.K.K., X.Z., and J.M.S. performed pathology and imaging analyses; S.A.K., L.M.P., and J.M.S. performed *in vitro* assays; and R.L.B. and J.W.H. wrote the paper with all the coauthors.

---

**Suppl. Fig. 1.**
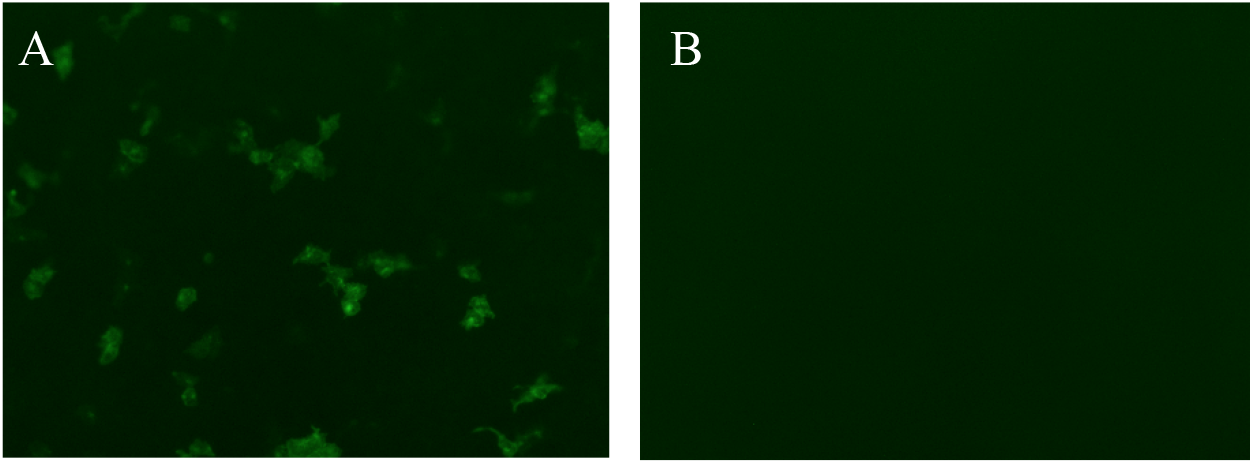
Expression of pWRG/nCoV-S(opt) plasmid in 293T cells. Expression of the spike protein was confirmed by transfection of 293T cells followed by immunofluroesence antibody test (IFAT) using **A)** human convalescent plasma and compared to **B)** empty vector.

**Suppl. Fig. 2.**
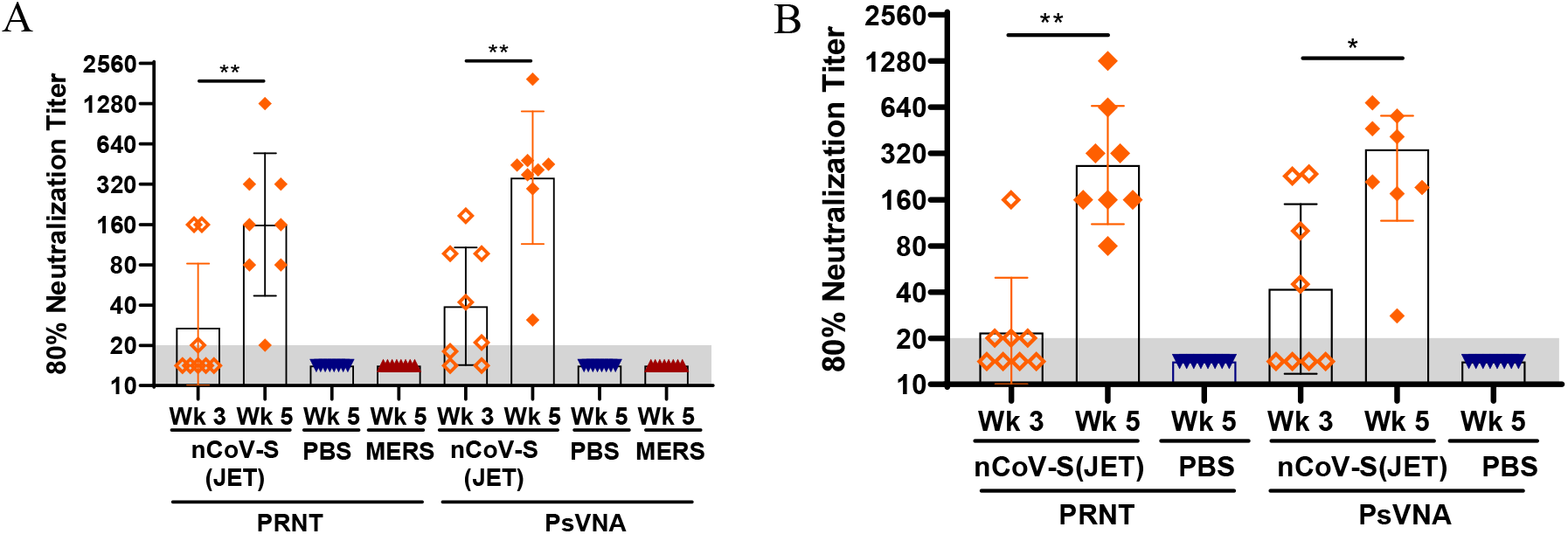
Neutralization titers of hamsters vaccinated with nCoV-S(JET) DNA vaccine. Eighty percent neutralization titers by PRNT and PsVNA of hamsters shown in **Figs. 1B** and **2B** are plotted.

**Suppl. Fig. 3.**
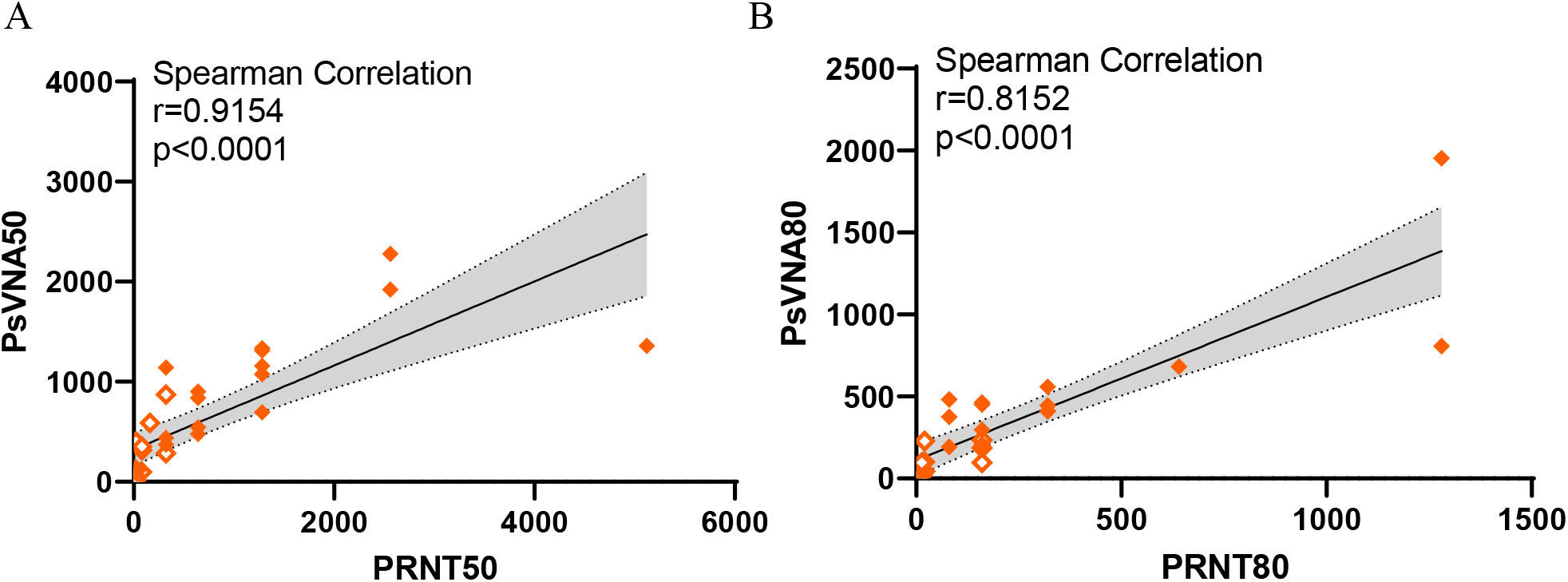
Correlation of PRNT and PsVNA assays. **A)** Fifty and **B)** 80 percent neutralization titers by PRNT and PsVNA from hamsters vaccinated once (open symbols) or twice (closed symbols) with the nCoV-S(JET) vaccine from **Fig. 1** and **Fig. 2** are plotted (positive by at least one assay only). Correlation was analyzed by Spearman with an **A)** r=0.9154 and *P<*0.0001 and **B)** r=0.8152 and *P*<0.0001 with linear regression (black line) and 95% confidence intervals (shaded area) shown.

**Suppl. Fig. 4.**
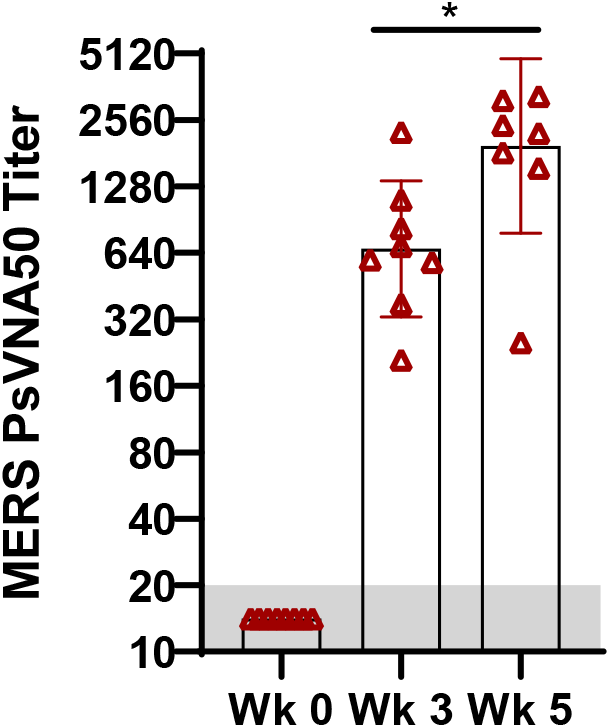
Neutralization titers of hamsters vaccinated with MERS-CoV DNA vaccine. Eight hamsters were vaccinated with MERS-CoV DNA vaccine using Tropis. Serum collected prior to and after 1 and 2 vaccinations was analyzed by MERS PsVNA (LLOQ = 20, grey shade). Asterisks indicate that results were statistically significant, as follows: *, *P*<0.05.

**Suppl. Fig. 5.**
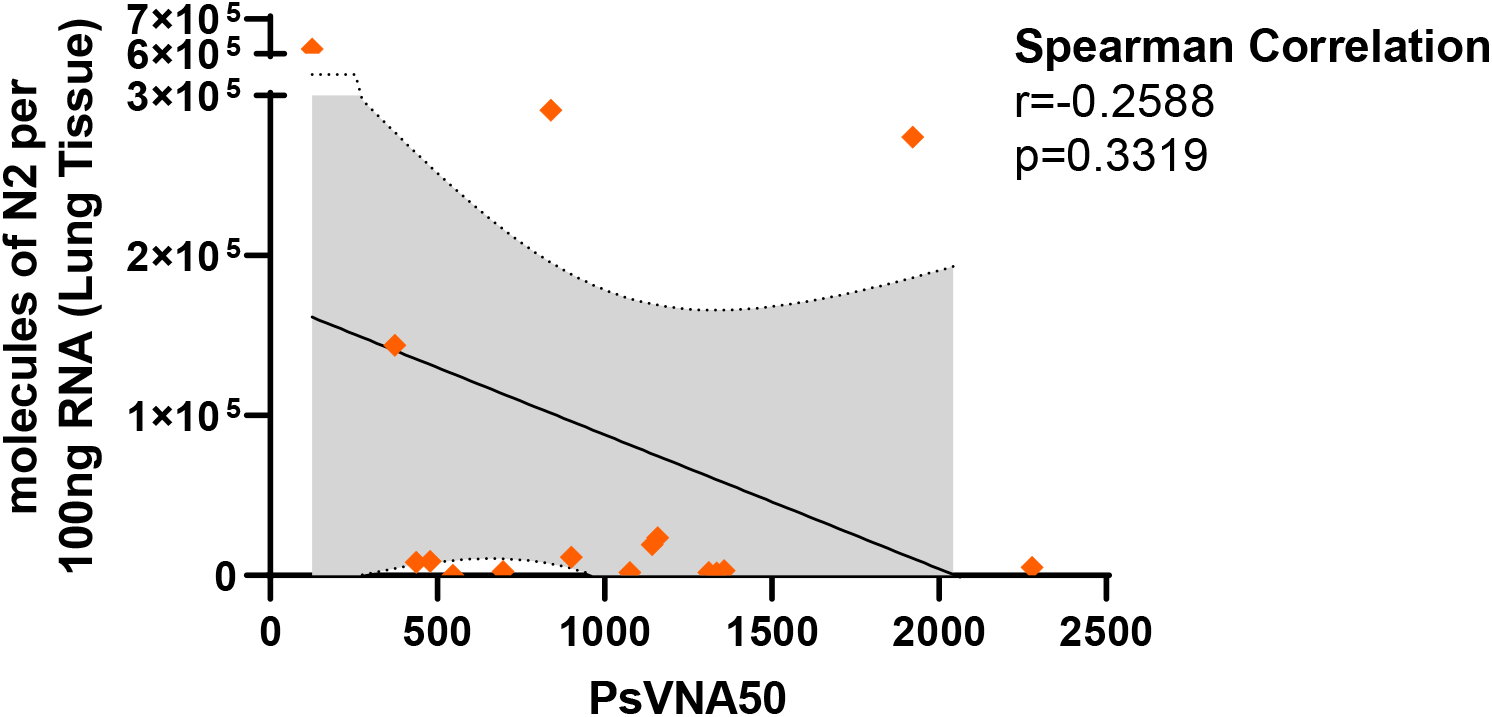
Correlation of PsVNA50 titers and lung viral RNA. Fifty percent neutralization titers by PsVNA from hamsters vaccinated twice with nCoV-S(JET) vaccine are plotted against viral RNA detected in lung homogenates at the time of euthanasia (both **Fig. 1** and **Fig. 2**). Correlation was analyzed by Spearman with an r=-0.2588 and *P*=0.3319 with linear regression (black line) and 95% confidence intervals (shaded area) shown.

